# Common dandelion (*Taraxacum officinale*) efficiently blocks the interaction between ACE2 cell surface receptor and SARS-CoV-2 spike protein D614, mutants D614G, N501Y, K417N and E484K *in vitro*

**DOI:** 10.1101/2021.03.19.435959

**Authors:** Hoai Thi Thu Tran, Nguyen Phan Khoi Le, Michael Gigl, Corinna Dawid, Evelyn Lamy

## Abstract

On 11th March 2020, coronavirus disease 2019 (COVID-19), caused by the SARS-CoV-2 virus, was declared as a global pandemic by the World Health Organization (WHO). To date, there are rapidly spreading new “variants of concern” of SARS-CoV-2, the United Kingdom (B.1.1.7), the South African (B.1.351) or Brasilian (P.1) variant. All of them contain multiple mutations in the ACE2 receptor recognition site of the spike protein, compared to the original Wuhan sequence, which is of great concern, because of their potential for immune escape. Here we report on the efficacy of common dandelion (*Taraxacum officinale*) to block protein-protein interaction of spike S1 to the human ACE2 cell surface receptor. This could be shown for the original spike D614, but also for its mutant forms (D614G, N501Y, and mix of K417N, E484K, N501Y) in human HEK293-hACE2 kidney and A549-hACE2-TMPRSS2 lung cells. High molecular weight compounds in the water-based extract account for this effect. Infection of the lung cells using SARS-CoV-2 spike pseudotyped lentivirus particles was efficiently prevented by the extract and so was virus-triggered pro-inflammatory interleukin 6 secretion. Modern herbal monographs consider the usage of this medicinal plant as safe. Thus, the *in vitro* results reported here should encourage further research on the clinical relevance and applicability of the extract as prevention strategy for SARS-CoV-2 infection.

**Significance statement:** SARS-CoV-2 is steadily mutating during continuous transmission among humans. This might eventually lead the virus into evading existing therapeutic and prophylactic approaches aimed at the viral spike. We found effective inhibition of protein-protein interaction between the human virus cell entry receptor ACE2 and SARS-CoV-2 spike, including five relevant mutations, by water-based common dandelion (*Taraxacum officinale*) extracts. This was shown *in vitro* using human kidney (HEK293) and lung (A549) cells, overexpressing the ACE2 and ACE2/TMPRSS2 protein, respectively. Infection of the lung cells using SARS-CoV-2 pseudotyped lentivirus was efficiently prevented by the extract. The results deserve more in-depth analysis of dandelions’ effectiveness in SARS-CoV-2 prevention and now require confirmatory clinical evidence.

## Introduction

In late 2019, the disease known as Corona Virus Disease 2019 or COVID-19 was first reported (1, 2). It is induced by the severe acute respiratory syndrome coronavirus 2 (SARS-CoV-2). Dry cough, fever, fatigue, headache, myalgias, and diarrhea are common symptoms of the disease. In severe cases people may become critically ill with acute respiratory distress syndrome (3). The SARS-CoV-2 virus surface is covered by a large number of glycosylated S proteins, which consist of two subunits, S1 and S2. The S1 subunit recognizes and attaches to the membrane-anchored carboxypeptidase angiotensin-converting enzyme 2 (ACE2) receptor on the host cell surface through its receptor binding domain (RBD). The S2 subunit plays a key role in mediating virus–cell fusion and in concert with the host transmembrane protease serine subtype 2 (TMPRSS2), promotes cellular entry (4). This interaction between the virus and host cell at entry site is crucial for disease onset and progression.

To date there are three rapidly spreading new variants of SARS-CoV-2 which were first reported in the United Kingdom (variant B.1.1.7), South Africa (variant B.1.351) and Brasil (variant P.1), all of which share the mutation N501Y in the spike protein (5). SARS-CoV-2 variants with spike protein D614G mutations now predominate globally. B.1.351 contains, besides D614G, other spike mutations, including three mutations (K417N, E484K und N501Y) in the RBD (6). Preliminary data suggest a possible association between the observed increased fatality rate with the mutation D614G and it is hypothesized that a conformational change in the spike protein results in increased infectivity (7). Free energy perturbation calculations for interactions of the N501Y and K417N mutations with both the ACE2 receptor and an antibody derived from COVID-19 patients raise important questions about the possible human immune response and the success of already available vaccines (8). Further, increased resistance of the variants B.1.351 and B.1.1.7 to antibody neutralization has been reported; for B.1.351 this was largely due to the E484K mutation in the spike protein (9).

Interference with the interaction site between the spike S1 subunit and ACE2 has the potential to be a major target for therapy or prevention (10). Compounds from natural origin may offer here some protection against viral cell entry while have no or few side effects. Here we report on the inhibitory potential of dandelion on the binding of the spike S1 protein RBD to the hACE2 cell surface receptor and compared the effect of the original D614 spike protein to its D614G, N501Y, and mix (K417N, E484K, N501Y) mutations.

The common dandelion (*Taraxacum officinale*) belongs to the plant family *Asteraceae*, subfamily *Cichorioideae* with many varieties and microspecies. It is a perennial herb, native distributed in the warmer temperate zones of the Northern Hemisphere inhabiting fields, roadsides and ruderal sites. *T. officinale* is consumed as vegetable food, but also employed in European phytotherapy to treat disorders from the liver, gallbladder, digestive tract or rheumatic diseases. Modern herbal monographs consider the plant usage as safe and have evaluated the empiric use of *T. officinale* with a positive outcome. Therapeutic indications for the use of *T. officinale* are listed in the German Commission E, the European Scientific Cooperative for Phytotherapy (ESCOP) monographs (11, 12) as well as in the British Herbal Medicine Association (13). The plant contains a wide array of phytochemicals including terpenes (sesquiterpene lactones such as taraxinic acid and triterpenes), phenolic compounds (phenolic acids, flavonoids, and coumarins) and also polysaccharides (14). The predominant phenolic compound was found to be chicoric acid (dicaffeoyltartaric acid). The other were mono-and dicaffeoylquinic acids, tartaric acid derivatives, flavone and flavonol glycosides. The roots, in addition to these compound classes, contain high amounts of inulin (15). Dosage forms include aqueous decoction and infusion, expressed juice of fresh plant, hydroalcoholic tincture as well as coated tablets from dried extracts applied as monopreparations (16) but also integral components of pharmaceutical remedies. Our research was conducted using water-based extracts from plant leaves. We found that leaf extracts efficiently blocked spike protein or its mutant forms to the ACE2 receptor, used in either pre-or post-incubation, and that high molecular weight compounds account for this effect. A plant from the same tribe (*Cichorium intybus*) could exert similar effects but with less potency. Infection of A549-hACE2-TMPRSS2 human lung cells using SARS-CoV-2 pseudotyped lentivirus was efficiently prevented by the extract.

## Results

### *T. officinale* inhibits spike S1 – ACE2 binding

We first investigated the inhibition of interaction between SARS-CoV-2 spike protein RBD and ACE2 using extracts from *T. officinale* leaves. In figure 1A, the concentration dependent inhibition of Spike S1 – ACE2 binding upon treatment with *T. officinale* extract is given (EC50=12 mg/ml). Extracts from *C. intybus*, also showed a concentration dependent binding inhibition, but with less potency than *T. officinale* (EC50=30 mg/ml) (figure 1B). We then prepared two fractions of the dried *T. officinale* as well as chicory leaves, separating the extracts into a high molecular (>5kDa) and low molecular weight (<5kDa) fraction. As can be seen from figure 1C, the bioactive compounds were mostly present in the HMW fraction. Only minor activity was seen in the LMW fraction.

**Figure 1:**
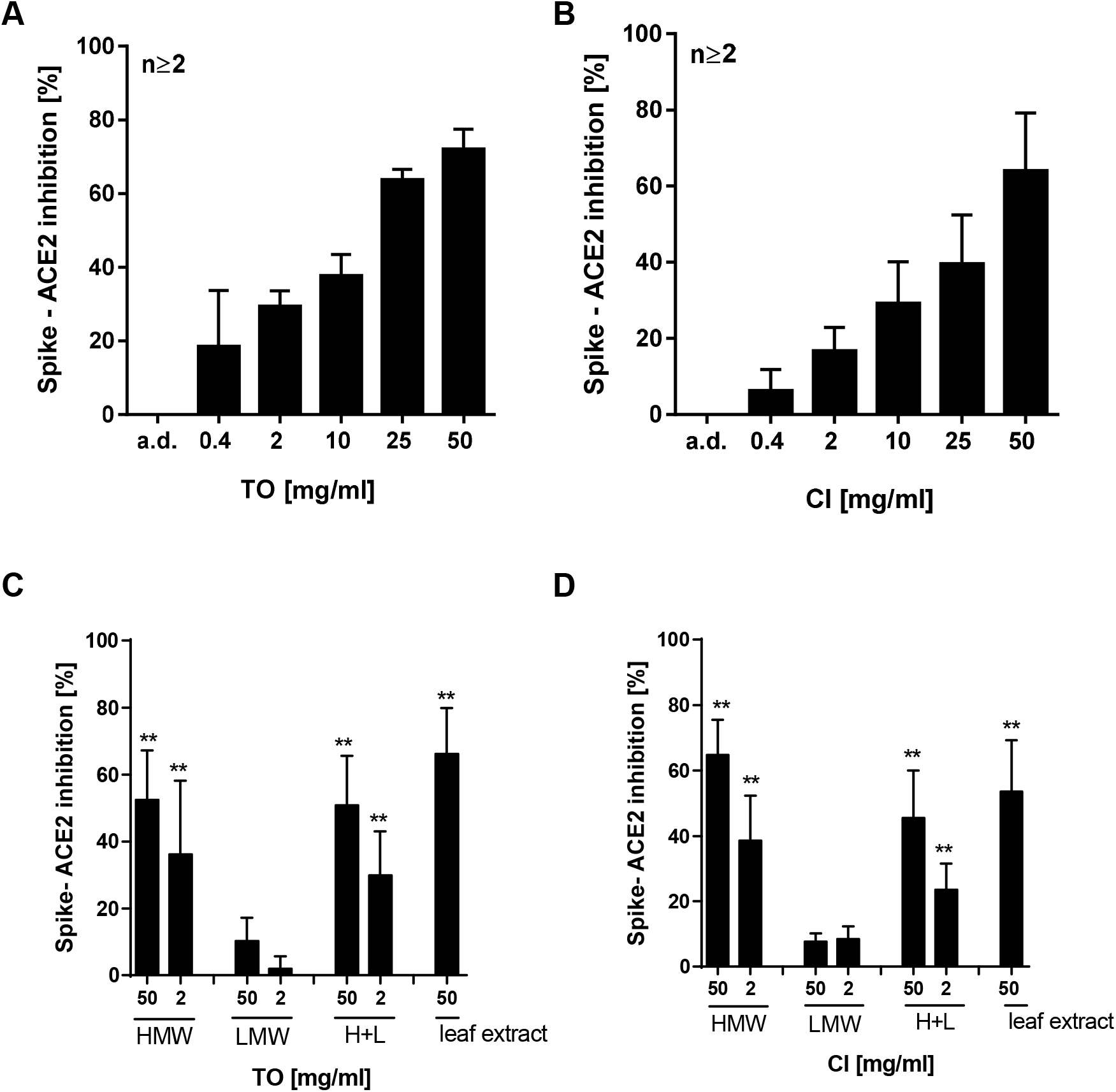
Effect of *T. officinale* and chicory on Sars-CoV-2-Spike – ACE 2 inhibition. A-B) Concentration dependent effect of *T. officinale* (TO) and *C. intybus* (CI) extract. C-D) Effect of fractions from TO and CI leave extract. The extracts were freeze-dried and a molecular weight fractionation subsequently carried out. The cut-off was set to 5 kDa (HMW > 5 kDa, LMW <5kDa). H+L: HMW and LMW fractions; 50 mg of dried leaves per ml water was used as reference. HMW and LMW fraction quantities equivalent to dried leaves were used. The binding inhibition was assessed using ELISA technique. Bars are means + SD. Solvent control: distilled water (a.d.).

Using hACE2 overexpressing HEK293 cells, the potential of *T. officinale* and *C. intybus* extracts to block spike binding to cells was further investigated. As can be seen from figure 2, pre-incubation of cells with *T. officinale* for one min. efficiently blocked cell binding of spike by 76.67% ± 2.9, and its HMW fraction by 62.5 ± 13.4% as compared to water control. After 3h, inhibition was still at 50 ± 13.6% for the extract, and 35.0 ± 20% for the HMW fraction of *T. officinale*. The chicory extract was less potent in this test system; binding inhibition was seen at 37 ± 20% after 1min. and 5.6 ± 9.9%.

**Figure 2:**
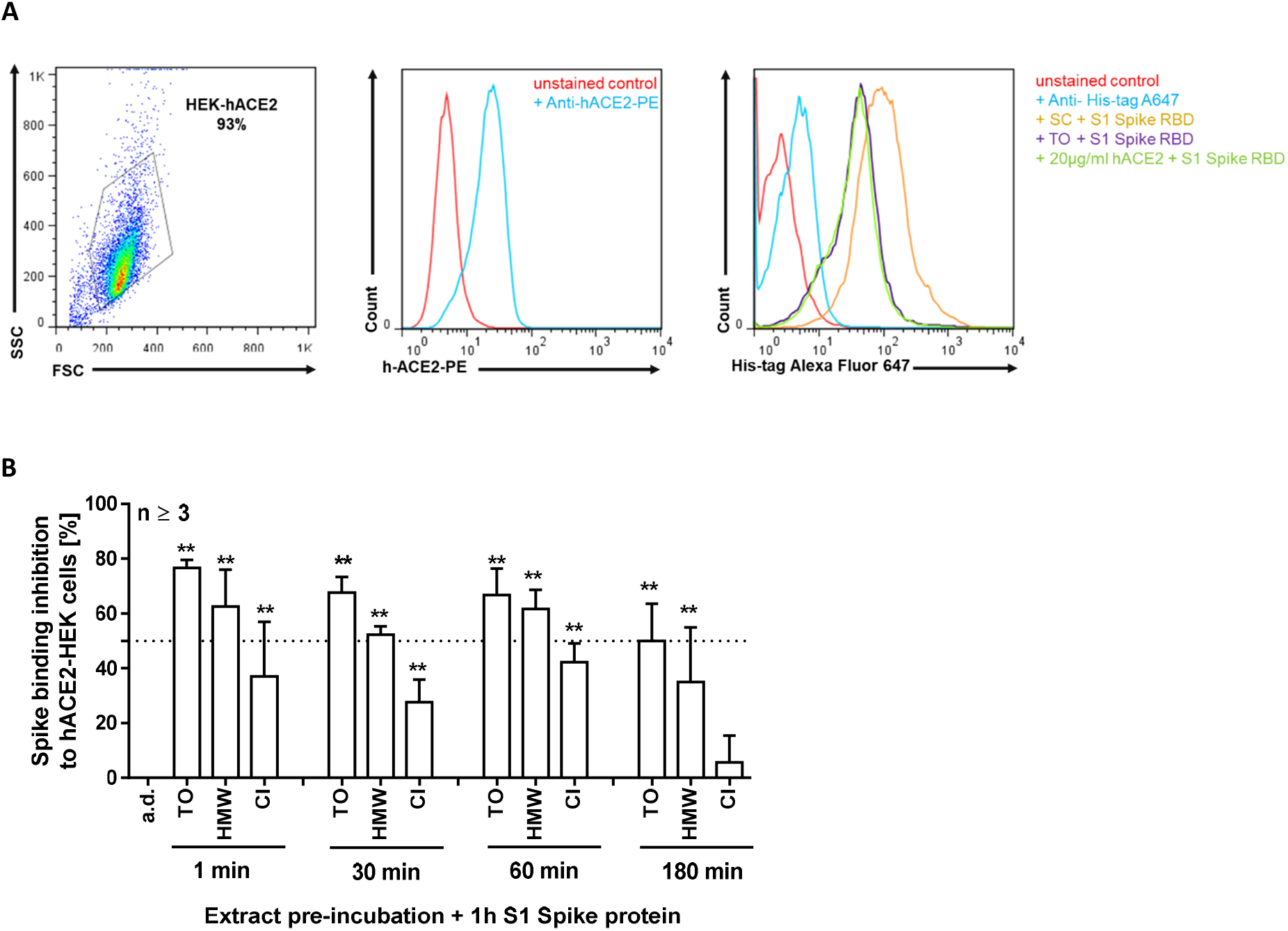
Binding inhibition of S1 spike protein to human HEK293-hACE2 cells by extract pre-incubation. Cells were pre-incubated for the indicated times with the extract from 10 mg/ml *T. officinale* (TO), its HMW fraction, equal to 10 mg/ml extract (HMW), and 10 mg/ml *C. intybus* (CI) or solvent control (a.d.) and subsequently treated with HIS-tagged S1 spike protein for 1h without a washing step in between at 4°C. Binding inhibition was assessed using flow cytometry. N=3, bars are means + SD. Upper left: cytogram of gated HEK-hACE2 cells. Middle: overlay of representative fluorescence intensity histograms for ACE2 surface expression. Upper right: overlay of representative fluorescence intensity histograms for spike binding inhibition by the extracts or a.d.; positive control: 20 µg/ml soluble hACE2. Cells were stained with anti-His-tag Alexa Fluor 647 conjugated monoclonal antibody.

Cell treatment with equal amounts of spike D614 and its variants D614G and N501Y confirmed a stronger binding affinity of D614G (about 1.5-fold) and N501Y (about 3 to 4-fold) than D614 spike protein to the ACE2 surface receptor of HEK293 cells (figure 3A). Pre-treatment with *T. officinale* quickly (within 30 sec.) blocked spike binding to the ACE2 surface receptor (figure 3B-C). After 30 sec., this was 58.2 ± 28.7% for D614, 88.2 ± 4.6% for D614G, and 88 ± 1.3% for N501Y binding inhibition by *T. officinale* extract. Even though for *C. intybus* extract a binding inhibition of spike could be seen, this was about 30-70% less compared to *T. officinale*, dependent on the spike protein investigated. When binding was studied at 37°C instead of 4°C, the results were comparable for *T. officinale*, but even weaker for chicory extract in this cell line (figure 3D). For *T. officinale* and *C. intybus* extracts the inhibition of spike binding was 47.90 ± 14.72 and 13.12 ± 12.37 (D614), 68.42 ± 14.53 and 8.86 ± 15.29 (D614G), 71.66 ± 7.66 and 37.56 ± 16.14 (N501Y), respectively. We also raised the question whether the extracts could replace spike binding to the ACE2 surface receptor of human cells. For this, we first incubated the cells with D614, D614G or N501Y spike protein and subsequently with the extracts. As given in figure 3D, *T. officinale* could potently remove spike from the receptor (on average 50%); chicory was much weaker then (on average 25%). We extended our experiments to human A549-hACE2-TMPRSS2 cells and could confirm the results observed in HEK293-hACE2 cells for *T. officinale* (figure 3D-G). This cell line has been stably transfected with both, the human ACE2 and TMPRSS2 genes and interestingly, here the *C. intybus* extract was more effective as compared to HEK-hACE2 cells. Upon extract pre-treatment, spike binding inhibition to the cells was between 73.5%± 5.2 (D614) to 86.3% ± 3.23 (N501Y) for *T. officinale* extract and 56.1% ± 5.28 (D614) to 63.07% ± 14.55 (N501Y) for *C. intybus* extract. Already at 0.6 mg/ml, *T. officinale* significantly blocked binding to D614G spike protein by about 40% (IC50 = 1.73 mg/ml). When cells were pre-incubated with the spike protein before extract treatment, results were comparable for *T. officinale* extract for D614 and D614G but somewhat lower for N501Y (figure 3C-D). Also in this settings, a mixture of spike mutants N501Y, K417N and E484K was tested and here again, *T. officinale* extract blocked binding by 82.97% ± 6.31(extract pre-incubation) and 79.7% ± 9.15 (extract post-incubation).

**Figure 3:**
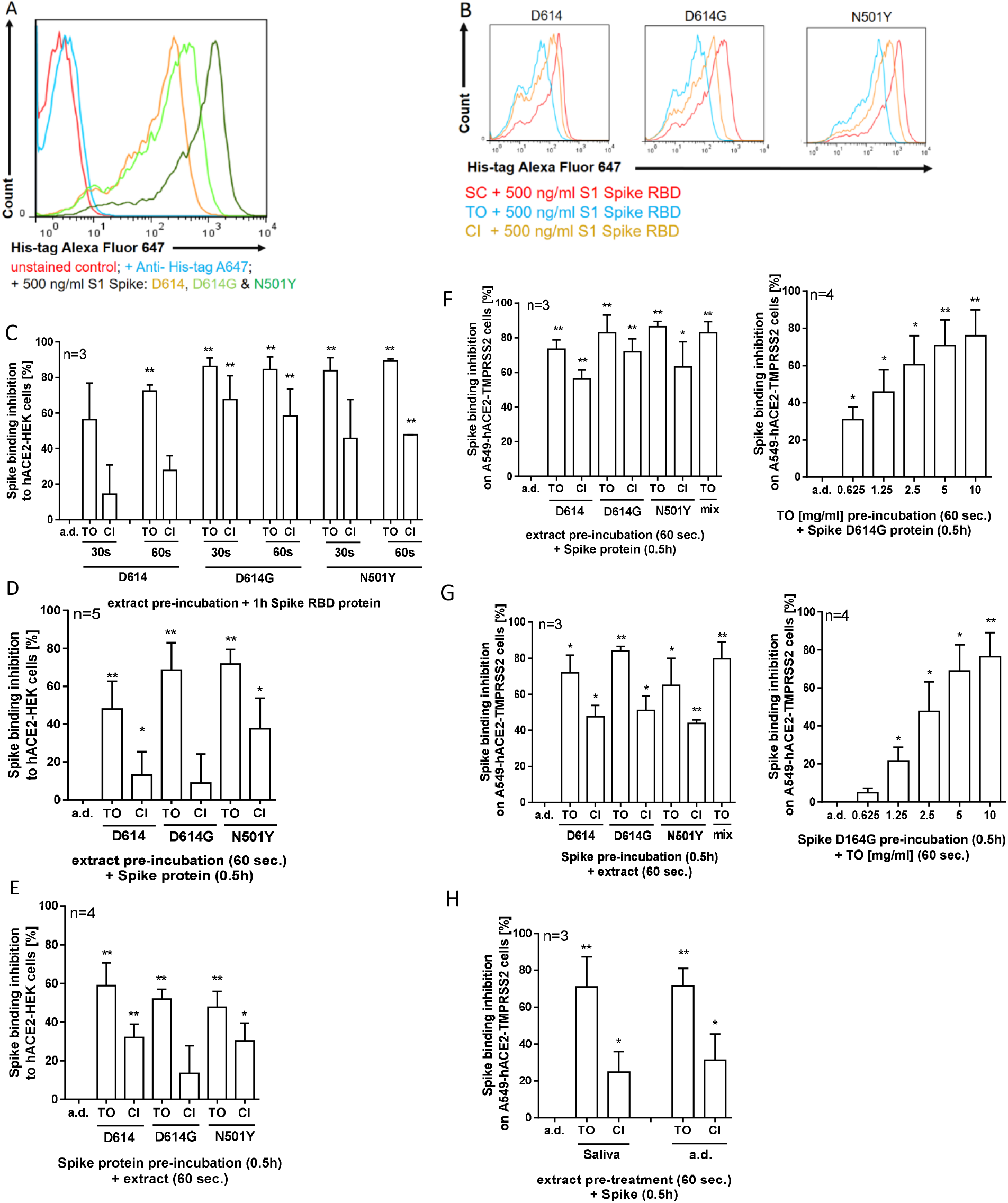
Binding inhibition of spike D614, and its mutants D614G, N501Y or mix (N501Y, K417N and E484K) to human HEK293-hACE2 and A549-hACE2-TMPRSS2 cells by extract pre- or post-incubation. Overlay of fluorescence intensity histogram for A) unstained HEK cells, staining control (anti-His-tag A647), and cells incubated with His-tag labelled spike D614, D614G or N501Y for 1h at 4°C. B) cells pre-incubated with solvent control (a.d.), 10 mg/ml *T. officinale* (TO) or 10 mg/ml *C. intybus* (CI) for 30-60 sec., and then treated with His-tag labelled S1 spike D614, D614G or N501Y protein for 1h without a washing step in between at 4°C. D-G) Effect of extract incubation on HEK or A549 cells either before or after incubation with His-tag labelled spike D614, D614G, N501Y or mix (N501Y, K417N and E484K) protein at 37°C. H) Plant extracts were incubated in saliva from 4 human donors for 30 min. at 37°C. Afterwards, cells were pre-treated with 5 mg/ml extracts for 60 sec. at 37°C before incubation with His-tag labelled spike D614 protein for 0.5h at 37°C. Spike binding inhibition to human cells was assessed using flow cytometric analysis of cells stained with anti-His-tag Alexa Fluor 647 conjugated monoclonal antibody. Bars are means +SD.

Extracts, incubated in human saliva for 30 min at 37°C before cell treatment had comparable effects on spike D614G inhibition (figure 3H) indicating a good stability of the bioactive compounds in saliva.

To see whether *T. officinale* extract interferes with the catalytic activity of the ACE2 receptor or affects ACE2 protein expression, we treated A549-hACE2-TMPRSS2 cells with the extract for 1-24h before cell lysis and detection. No loss in cell viability was seen after extract exposure to the cells for 84h (figure 4A). No enzyme activity impairment could be detected after 1 or 24h (4B). Spike significantly downregulated ACE2 protein after 6h (4C, black bars), and this was also true for the extract, either alone (4C, white bars) or in combination with spike (black bars). After 24h, this effect was abolished (4D).

**Figure 4:**
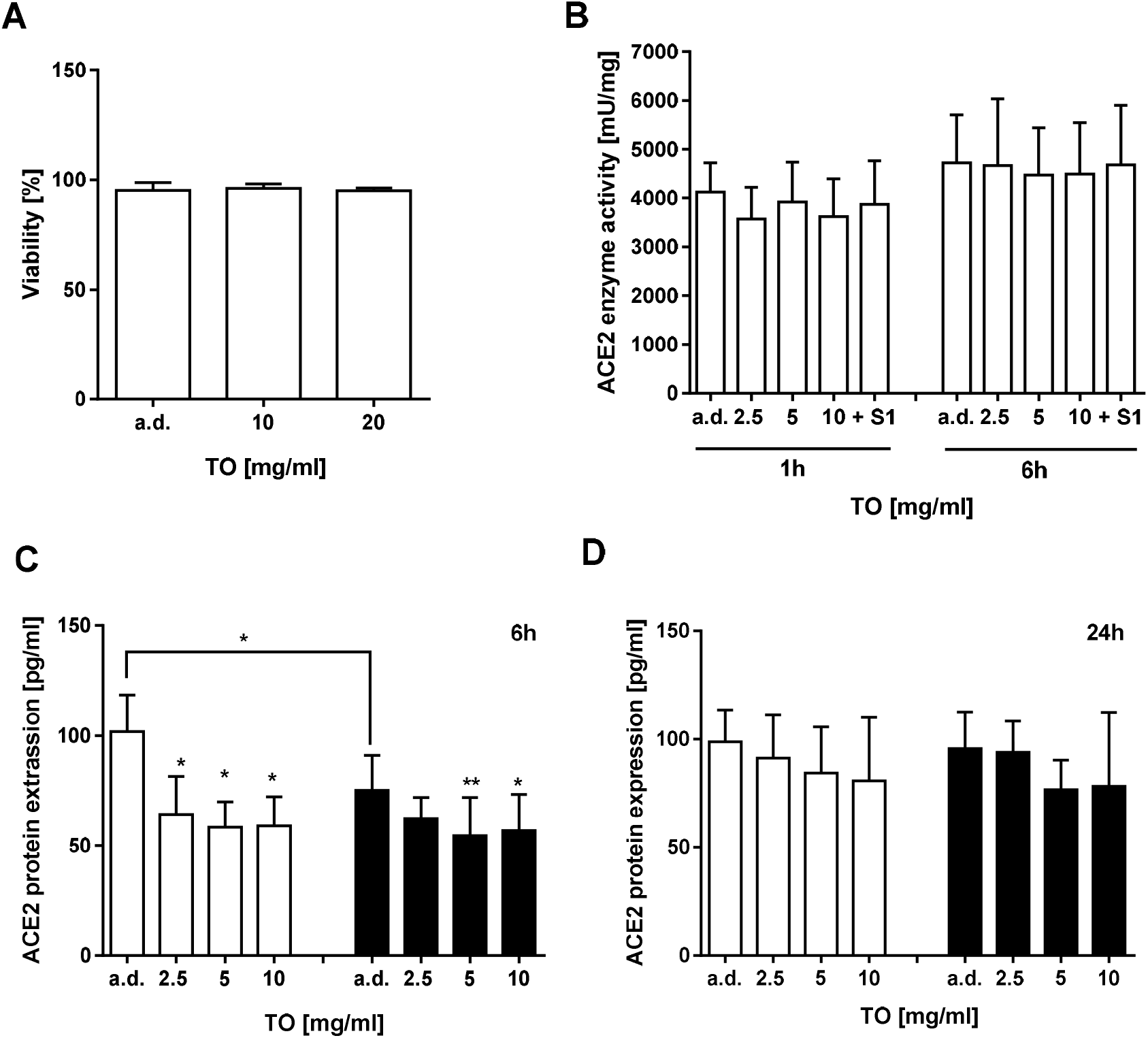
Effect of *T. officinale* extract on ACE2 enzyme activity and protein expression. A) Viability of A549-hACE2-TMPRSS2 cells was determined using trypan blue cell staining after 84h exposure to the extract. B) Cells were incubated with TO extract or 500 ng/ml S1 protein and analysed for enzyme activity using a fluorescence kit. C-D) Cells were exposed for 6h or 24h to extract without (white bars) or with (black bars) 500 ng/ml S1 protein and analysed for ACE2 protein expression using a human ACE2 ELISA kit; a. d.: solvent control. Bars are means + SD, N ≥ 3 independent experiments.

Using a SARS-CoV-2 spike pseudotyped lentivirus, we then studied whether the extract could block virus entry via spike inhibition. When pre-treated with the extract, virus transduction was diminished by about 85% at 20 mg/ml (figure 5A). Under the different treatment conditions, the luminescent signal produced by virus transduction was inhibited at 10 mg/ml extract by 70% ± 16.7 (A), 58% ± 9.6 (B) and 53% ± 8.1 (C). This inhibition of virus transduction by the extract concurred with a significant suppression of the virus triggered inflammatory response, as determined by diminished secretion of the pro-inflammatory cytokine IL-6 in A549-hACE2-TMPRSS2 cells (figure 5D).

**Figure 5.**
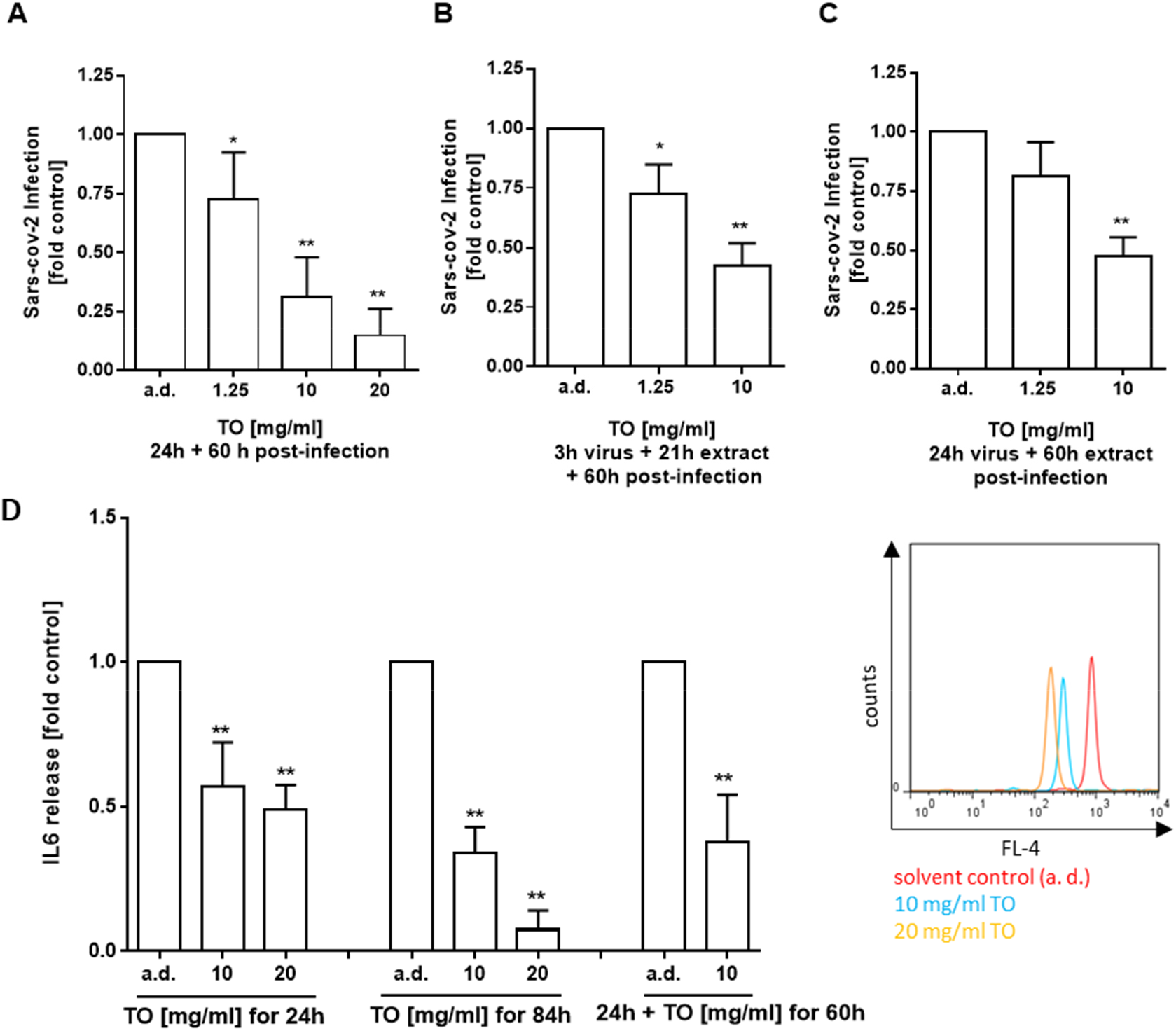
Viral transduction inhibition of A549-hACE2-TMPRSS2 cells by *T. officinale* extract. Cells were transduced with 2.5µl SARS-CoV-2-Spike pseudotyped lentivirus (Luc reporter) for 24h A) after pre-treatment with *T. officinale* (TO) extract for 0.5 h, B) 3h before addition of TO or C) without extract. Afterwards, the medium was changed to fresh medium and cells incubated for another 60h together with the extract. Luminescence was detected after 1h. (-) negative control: bald lentiviral pseudovirion; (+) positive control: firefly luciferase lentivirus. D) pro-inflammatory IL-6 cytokine secretion analysis was done either after 24h virus transduction together with extract (left), after 24h + 60h post-infection with extract (middle) or after 60h post-infection with extract (right) using multiplexing flow cytometric analysis. Solvent control: destilled water (a.d.). N ≥ 3 independent experiments.

## Discussion

The development of effective prevention and treatment strategies for SARS-CoV-2 infection is still at the early stages. Although the first vaccines received now marketing authorization, challenges in distribution concerns or about durable effectiveness, and risk of re-infection remain (17, 18). Subsequent infections may possibly be milder than the first one, though. Besides vaccination against COVID-19, blocking the accessibility of the virus to membrane-bound ACE2 as the primary receptor for SARS-CoV-2 target cell entry, represents an alternative strategy to prevent COVID-19. Here, different approaches exist (19), but of course each of these treatment strategies also has its fundamental as well as translational challenges which need to be overcome for clinical utility. Technical hurdles include off-target potential, ACE2-independent effects, stability or toxicity (19). Compounds from natural origin could be an important resource here as they are described long term and many of them have been established as safe. While *in silico* docking experiments suggested different common natural compounds as ACE2 inhibitors, spike binding inhibition to ACE2 has not been shown for most of them so far, which might be explained by a lack of complete coverage of ACE2 binding residues by the compounds (20). However, for glycyrrhizin, nobiletin, and neohesperidin, ACE2 binding falls partially within the RBD contact region and thus, these have been proposed to additionally block spike binding to ACE2 (20). The same accounts for synthetic ACE2 inhibitors, such as N-(2-aminoethyl)-1 aziridine-ethanamine (NAAE) (21). In contrast, the lipoglycopeptide antibiotic dalbavancin has now been identified as both, ACE2 binder and SARS-CoV-2 spike-ACE2 inhibitor (22); SARS-CoV-2 infection was effectively inhibited in both mouse and rhesus macaque models by this compound. Also, for a hydroalcoholic pomegranate peel extract, blocking of spike-ACE2 interaction was shown at 74%, for its main constituents punicalagin at 64%, and ellagic acid at 36%. Using SARS-CoV-2 spike pseudotyped lentivirus infection of human kidney-2 (HK-2) cells, virus entry was then efficiently blocked by the peel extract (23). In the present study, we could show potent ACE2-spike S1 RBD protein inhibition by *T. officinale* extracts using a cell-free assay and confirmed this finding by demonstrating efficient ACE2 cell surface binding inhibition in two human cell lines. We observed stronger binding of the variants D614G and N501Y to the ACE2 surface receptor of human cells, but all tested variants were sensitive to binding inhibition by *T. officinale*, either used before spike protein exposure or thereafter. To date, several studies indicate that the D614G viral lineage is more infectious than the D614 virus (24). Also, the presence of characteristic mutations such as N501Y of, e. g. the so-called UK variant B.1.1.7, result in higher infectivity than the parent strain which might be due to a higher binding affinity between the spike protein and ACE2 (25). So our findings on *T. officinale* extracts could here be important, as with progression of the pandemic, new virus variants of potential concern will emerge which may also reduce the efficacy of some vaccines or cause increased rates of reinfections. As mentioned above, an issue in the development of products as prophylaxis for SARS-CoV-2 infection or for slowing the systemic virus spread, is the selectivity towards virus intrusion with low toxicity needed for the host. For current medical indications, no case of overdose by *T. officinale* has been reported (11, 13, 16). The recommended dosage is 4–10 g (about 20-30 mg per ml hot water) up to 3 times per day (Commission E and ESCOP). Based on the information provided by the European Medicines Agency (EMA) contraindications for the use of *T. officinale* are hypersensitivity to the *Asteraceae* plant family or their active compounds, liver and biliary diseases, including bile duct obstruction, gallstones and cholangitis, or active peptic ulcer (16). The plant is a significant source of potassium (26, 27) and thus a warning is given because of the possible risk for hyperkalaemia. The use in children under 12 years of age, during pregnancy and lactation has not been established due to lack of adequate or sufficient data.

While ACE2 enzyme activity was not affected by *T. officinale* extract in the present study, ACE2 protein was transiently downregulated in the ACE2 overexpressing lung cell line, which requires more attention in ongoing studies. ACE2 is an important zinc-dependent mono-carboxypeptidase in the renin-angiotensin pathway, critical in impacting cardiovascular and immune system. Disruption of the Angiotensin II/Angiotensin-(1-7) balance by ACE2 enzyme activity inhibition or protein decrease and more circulating Angiotensin II in the system, is e. g. recognised to promote lung injury in the context of COVID-19 disease (28, 29).

The lung would be assumed to be the primary target of interest but, ACE2 mRNA and protein expression have been found in epithelial cells of all oral tissues, especially in the buccal mucosa, lip and tongue (30). These data concur with the observation of very high salivary viral load in SARS-CoV-2 infected patients (31, 32). As an essential part of the upper aerodigestive tract, the oral cavity is thus believed to play a key role in the transmission and pathogenicity of SARS-CoV-2. There is high potential that prevention of viral colonization at the oral and pharyngeal mucosa could be critical for averting further infection to other organs and the onset of COVID-19 (33). Commercial virucidal mouth-rinses, povidone-iodine at the first place, have thus been suggested to potentially reduce the SARS-CoV-2 virus load in infected persons (34-36), but significant clinical studies do not exist to date (36). Blocking SARS-CoV-2 virus binding to cells of the oral cavity with *T. officinale* extracts might be tolerable for a consumer, if necessary only for limited periods of time (e. g. product application after contact with infected persons or when being infected). More physiologically relevant *in vitro* experiments that were carried out by us showed that only short contact times with *T. officinale* extract were necessary for efficient blocking of SARS-CoV-2 spike binding or for removing already bound spike from the cell surface. Further evidence of relevance was given using SARS-CoV-2 spike pseudotyped virus experiments. Even though the use of these pseudotyped viruses does not allow to assess the contribution of virion characteristics, such as membrane or envelope proteins, on the cell tropism (37), they are regarded as a useful tool to document the relevance of ACE2 for the cell entry steps mediated by the spike protein.

Developed vaccine candidates all aim to generate antibody (and T cell) responses against the spike protein and spike sequences from the early Wuhan strain served here as basis (38). However, SARS-CoV-2 is steadily mutating during continuous transmission among humans. Virus antigenic drift is clearly shown by the recent appearance of B.1.1.7, B.1.351, or B.1.1.28 (P.1). It is evolving in such a way that it may eventually be able to evade our existing therapeutic and prophylactic approaches aimed at the viral spike. Thus, factors such as low toxicity in humans and effective binding inhibition of five relevant spike mutations to the human ACE2 receptor, as reported here *in vitro*, encourage for more in-depth analysis of *T. officinales*’ effectiveness in SARS-CoV-2 prevention and now requires further confirmatory clinical evidence.

## Material and Methods

### Plant Material

The study was carried out using dried leaves from *T. officinale* (vom Achterhof, Uplengen, Germany; batch no. 37259, B370244, and P351756). Plant leaf samples were also collected at three different places in the region of Freiburg i. Br. (Germany), on 12.7.2020, and tested positive in the cell-free spike S1-ACE2 binding assay (data not shown). *C. intybus* was purchased from Naturideen (Germany).

### Cell lines and culture conditions

Human embryonic kidney 293 (HEK293) cells, stably expressing hACE2, were generously provided by Prof. Dr. Stefan Pöhlmann (Göttingen, Germany). The cells were maintained in Dulbecco’s modified Eagle medium (DMEM), high glucose supplemented with 10% fetal calf serum (FCS), 100 U/ml penicillin/streptomycin and 50 µg/ml zeocin (Life Technologies, Darmstadt, Germany). Human A549-hACE2-TMPRSS2 cells, generated from the human lung A549 cell line were purchased from InvivoGen SAS (Toulouse Cedex 4, France) and maintained in DMEM, high glucose supplemented with 10% heat-inactivated FCS, 100 U/ml penicillin/streptomycin, 100 µg/ml normocin, 0.5 µg/ml puromycin and 300 µg/ml hygromycin. To subculture, all cells were first rinsed with phosphate buffered saline (PBS) then incubated with 0.25% trypsin-EDTA until detachment. All cells were cultured at 37 °C in a humidified incubator with 5% CO_2_/95% air atmosphere.

### Plant extraction

Dried plant material was weighted in an amber glass vial (Carl Roth GmbH, Germany) and mixed with HPLC-grade water (a.d.), at room temperature (RT). Extracts were then incubated for 1h and centrifuged at 16.000g (3 min, RT). The supernatant was filtered (0.22 µm) prior to use for the experiments.

### Analysis of SARS-COV2 Spike – ACE2 interaction inhibition using ELISA and flow cytometry

A commercially available SARS-CoV-2 Inhibitor Screening Kit (Cat#: 16605302, Fisher Scientific GmbH, Schwerte, Germany) was used for cell free detection of SARS-CoV-2 Spike – ACE2 interaction inhibition. This colorimetric ELISA assay measures the binding between immobilized SARS-CoV-2 spike protein RBD and biotinylated human ACE2 protein. The colorimetric detection is done using streptavidin-HRP followed by TMB incubation. A SARS-CoV-2 inhibitor (hACE2) was used as method verified reference.

Cell surface expression of ACE2 was determined by using a human ACE2 PE-conjugated antibody (Bio-Techne GmbH, Wiesbaden-Nordenstadt, Germany) and flow cytometric analysis. For analysis of SARS-CoV-2 S1 Spike RBD -ACE2 binding, 2×10^5^ cells (5×10^6^ cells/ml) were pre-treated with plant extracts for different time points. Then, 500 ng/ml SARS-CoV-2 Spike S1 (Trenzyme GmbH, Konstanz, Germany), spike S1 D614G, N50Y, or mix of K417N, E484K, and N501Y (Sino Biological Europe GmbH, Eschborn, Germany) -His recombinant protein were added into each sample, and samples were further incubated for 30-60 min. In another setting, cells were pre-treated with 500 ng/ml SARS-CoV-2 Spike -His recombinant protein for 30 min prior to incubation with the plant extract for 30-60 sec at 4°C or 37°C. The samples were incubated in PBS buffer containing 5% FCS. Cells were then washed one time with PBS buffer containing 1% FCS at 500 x g, 5 min before staining with His-tag A647 mAb (Bio-Techne GmbH, Wiesbaden-Nordenstadt, Germany) for 30 min at RT. Subsequently, cells were washed twice as described above. The cells were analysed by using a FACSCalibur (BD Biosciences, Heidelberg, Germany), 10.000 events were acquired. The median fluorescence intensity (MFI) of each sample were determined using FlowJo software (Ashland, Oregon, USA).

### Human ACE2 enzyme activity and protein quantification

A549-hACE2-TMPRSS2 (2×10^5^) cells were seeded in a 24-well plate in high glucose DMEM medium, containing 10% heat inactivated FCS, at 37°C, 5 % CO_2_. Cells were then treated with *T. officinale* extract with/without 500 ng/ml SARS-CoV-2 S1 Spike RBD protein for 1-24h. Afterwards, cells were washed with PBS and lysed. 25µg protein were used for quantification of ACE2 protein (ACE2 ELISA kit), 5 µg for ACE2 enzyme activity (ACE2 activity assay kit, Abcam, Cambridge, UK) according to the manufacturer’s instructions.

### Infection of A549-hACE2-TMPRSS2 cells using SARS-CoV-2 pseudotyped lentivirus

SARS-CoV-2 spike pseudotyped lentivirus particles, produced with SARS-CoV-2 spike (Genbank Accession #QHD43416.1) as the envelope glycoproteins instead of the commonly used VSV-G, were purchased from BPS Bioscience, (Catalog#: 79942, Biomol, Hamburg). These pseudovirions also contain the firefly luciferase gene driven by a CMV promoter. Thus, the spike-mediated cell entry can be quantified via luciferase reporter activity. The bald lentiviral pseudovirion (BPS Bioscience #79943), where no envelope glycoprotein is expressed, was used as a negative control. The Firefly Luciferase Lentivirus (Puromycin) from BPS Bioscience, (catalogue#: 79692-P) was used as positive control for transduction. These viruses constitutively express firefly luciferase under a CMV promoter. Lung cells were seeded at 0.1×10^6^ cells/cm2 in 96-well plate in DMEM containing 10% heat-inactivated FCS, 100 U/ml penicillin/streptomycin, 100 µg/ml normocin, 0.5 µg/ml puromycin and 300 µg/ml hygromycin overnight. The medium was replaced by DMEM + 10% heat-inactivated FCS and cells were treated with a.d. or *T. officinale* extract either 30 min. before or 3h after addition of 2.5 µl of the lentivirus particles. After 24h of virus particle incubation, the medium was removed by washing with PBS, fresh medium was added and cells incubated for another 60h with the addition of a.d. or *T. officinale* extract. Luminescence was detected within 1h using the one-step luciferase reagent from BPS following the manufacturer’s protocol in a multiplate reader from Tecan (Tecan Group Ltd, Crailsheim, Germany).

### Quantification of cytokine release by multiplex bead technique

After 24h SARS-CoV-2 spike pseudotyped lentivirus transduction and 60h post-infection of A549-hACE2-TMPRSS2 cells, supernatants were collected and stored at −80 °C until analysis for cytokine secretion using the human MACSplex cytokine 12-kit (Miltenyi Biotec GmbH, Bergisch Gladbach, Germany) according to manufacturer’s protocol.

### Molecular weight fractionation from plant extracts

Extracts from dried plant leaves were prepared by adding bidistilled water (5 ml) to plant material (500 mg each). The samples were incubated in the dark at room temperature (RT) for 60 min, followed by centrifugation at 16.000 g for 3 min. The supernatants were collected and membrane filtrated (0.45 µm), resulting in the extracts. Aliquots were freeze dried for 48h to determine their yield by weight. The extracts were then further separated in a high molecular weight (HMW) and low molecular weight (LMW) fraction, using a centrifugation tube with an insert containing a molecular weight cut-off filter (5 kDa, Sartorius Stedim Biotech, Goettingen, Germany). Each HMW fraction was purified by flushing with 20 ml of water, yielding the HMW fractions, as well as LMW. The fractions were freeze dried, their yield determined by weight and stored at −20°C until use.

### Determination of cell viability using trypan blue staining

Cell viability was assessed using the trypan blue dye exclusion test as described before (Odongo et al., 2017). Briefly, A549-hACE2-TMPRSS2 cells were cultured for 24h, and then exposed to extracts or the solvent control (a. d.) for 84 h.

### Statistical analysis

Results were analysed using the GraphPad Prism 6.0 software (La Jolla, California, USA). Data were presented as means + SD. Statistical significance was determined by the one-way ANOVA test followed by Bonferroni correction. P values <0.05 (*) were considered statistically significant and <0.01 (**) were considered highly statistically significant.

## Acknowledgements

The authors are grateful to Prof. Dr. Stefan Pöhlmann (German Primate Center, Göttingen, Germany) for providing the human embryonic kidney 293 (HEK293) cells, stably expressing hACE2.

## Authors contributions

Study design and conception: E.L.; design of experiments, data acquisition, analysis of data: H.T.T, E. L., N.P.K.L.; preparation of extract fractions: C.D., M.G.; writing of the first manuscript draft: E.L.. All authors commented on previous versions of the manuscript.

